# Eigenvalue Signatures Reveal Residual Motion Effects Across Resting-State fMRI Denoising Strategies

**DOI:** 10.64898/2026.09.08.750194

**Authors:** Lejian Huang, Andrew D. Vigotsky, A. Vania Apkarian

## Abstract

The eigenvalue structure of resting-state fMRI (RS-fMRI) signals provides a compact representation of its variance-covariance structure, yet the information it encodes remains unclear. In this study, we introduce an eigenvalue-based framework to characterize RS-fMRI data using two features derived from the eigenspectrum: the log_10_-transformed first eigenvalue (log_10_(*λ*_1_)) and the slope of the log_10_-transformed spectrum (*β*). We systematically evaluated these features across 14 denoising strategies using two independent datasets (China167 (83 males, 84 females; mean age ± SD: 41 ± 14 years old) and HCP1200 (425 males, 501 females; mean age ± SD: 29 ± 4 years old)). Specifically, we used singular value decomposition to obtain the eigenspectrum of the cortical BOLD signals after preprocessing and nuisance regression, and a linear model to parameterize it. We then examined the relationships between the eigenvalue parameters, head motion, and functional connectivity metrics across subjects and denoising strategies.

Our key findings include: (1) log_10_(*λ*_1_) and *β* strongly covaried with one another across subjects, denoising strategies, and datasets, indicating that they capture highly coherent aspects of the eigenspectrum; (2) both features were systematically influenced by denoising strategies; (3) within each denoising strategy, participants with greater log_10_(*λ*_1_) had greater mean framewise displacements (mFD), demonstrating sensitivity to residual motion effects, for all strategies in HCP1200 and 12/14 in China167; (4) across denoising strategies, mean log_10_(*λ*_1_) reflected the proportion of functional connectivity edges significantly associated with motion, indicating that higher log_10_(*λ*_1_) reflects more widespread motion-related contamination across large-scale functional networks; and (5) global signal regression consistently reduced log_10_(*λ*_1_), whereas spike regression had dataset-dependent effects.

Together, these results establish eigenvalue signatures as robust and sensitive metrics for quantifying residual motion effects in RS-fMRI and provide a framework for evaluating denoising performance based on eigenvalue structure.

The full procedure is implemented in R, and the corresponding script is available at: https://github.com/lejianhuang/EigenvalueSignature.

## 1. Introduction

In RS-fMRI studies, eigenvalue analysis, commonly implemented through principal component analysis (PCA), provides a compact spectral representation of RS-fMRI signals by decomposing the variance-covariance structure of the BOLD time series into orthogonal components ordered by their variance contributions (Jolliffe 2002, Leonardi, Richiardi et al. 2013). The leading eigenvalues and their corresponding eigenvectors capture dominant patterns of neural activity, such as brain network organization (Zajac and Piersa 2013, Wang, Owen et al. 2017, Bansal and Peterson 2021, Ghosh, Raj et al. 2024), as well as physiological processes, motion-related influences, and scanner-induced artifacts (Behzadi, Restom et al. 2007, Pruim, Mennes et al. 2015, Soltysik, Thomasson et al. 2015). Discarding noise-dominated components while preserving those that primarily capture the signal can facilitate effective RS-fMRI denoising (Behzadi, Restom et al. 2007, Zhu, Ma et al. 2022). However, doing so requires understanding the provenance of RS-fMRI eigenspectra. Here, we explore and demonstrate the sensitivity of the RS-fMRI eigenspectrum to residual motion effects and how this sensitivity critically depends on denoising strategies, highlighting that the covariance structure of the RS-fMRI signal can be leveraged to inform denoising.

Head motion during scanning degrades the quality of RS-fMRI BOLD signals by introducing non-neuronal fluctuations (Friston, Williams et al. 1996, Power, Barnes et al. 2012, Satterthwaite, Wolf et al. 2012, Van Dijk, Sabuncu et al. 2012). These motion effects are more pronounced in patient populations, such as those with chronic pain or Parkinson’s disease (Yang, Wu et al. 2021, Reddy, Zvolanek et al. 2024), and in those with high body mass index (BMI) (Huang, Vigotsky et al. 2024), highlighting the importance of rigorous preprocessing and denoising in these populations (Golestani and Chen 2022, Pavlovich, Pang et al. 2025). However, standard denoising approaches do not fully remove motion-related variance, as indicated by two quality-control (QC) metrics: residual QC–FC (QC–functional connectivity) correlations and dissimilarities in functional connectivity. Each of these captures the influence of residual motion effects across populations, scanners, and preprocessing pipelines (Power, Barnes et al. 2012, Ciric, Wolf et al. 2017, Yang, Wu et al. 2021). Although generally undesirable, the sensitivity of FC to motion artifacts makes FC-based metrics useful for choosing denoising strategies.

Because FC is simply the correlation matrix of RS-fMRI signals, motion artifacts that systematically alter FC should also perturb RS-fMRI eigenspectra. An eigenspectrum-based QC metric would be timely, as it could facilitate computationally efficient QC to accommodate the growing prevalence of large fMRI datasets. Specifically, researchers could forgo estimating large or rank-deficient correlation matrices and instead use singular value decomposition (SVD) to efficiently extract the eigenvalues of the top *k* components. However, before establishing such an eigenspectrum-based QC metric, it must be empirically validated. Thus, this study aimed to evaluate how residual motion effects depend on the denoising strategy and whether they can be detected through eigenvalue analysis.

## 2. Methods

### 2.1. Study Design

The overall study design is illustrated in **Fig. 1a** and consists of four sequential steps. First, **preprocessing**: all raw RS-fMRI data was preprocessed using the fMRIPrep software (Esteban, Markiewicz et al. 2019), including standard procedures such as motion correction, spatial normalization to MNI152 space, and the extraction of confounding variables (details are expounded below). Second, **denoising**: one of 14 denoising strategies was selected and combined with confounds obtained from the preprocessing step to construct a nuisance regressor matrix. We used this matrix to regress out confounding effects from the preprocessed RS-fMRI data, followed by temporal low-pass filtering with a cut-off frequency of 0.2 Hz. Third, **eigenvalue generation**: principal component analysis (PCA) via singular value decomposition was performed on *z*-scored voxel-wise BOLD time series within a common cortical mask derived from datasets used in the study. This procedure yielded eigenvalues that characterize the variance structure of cortical BOLD signals for each subject. Fourth, **analysis**: eigenvalue-derived features were modeled, and their properties and relationships with the strategies and residual motion were systematically evaluated.

**Figure 1.**
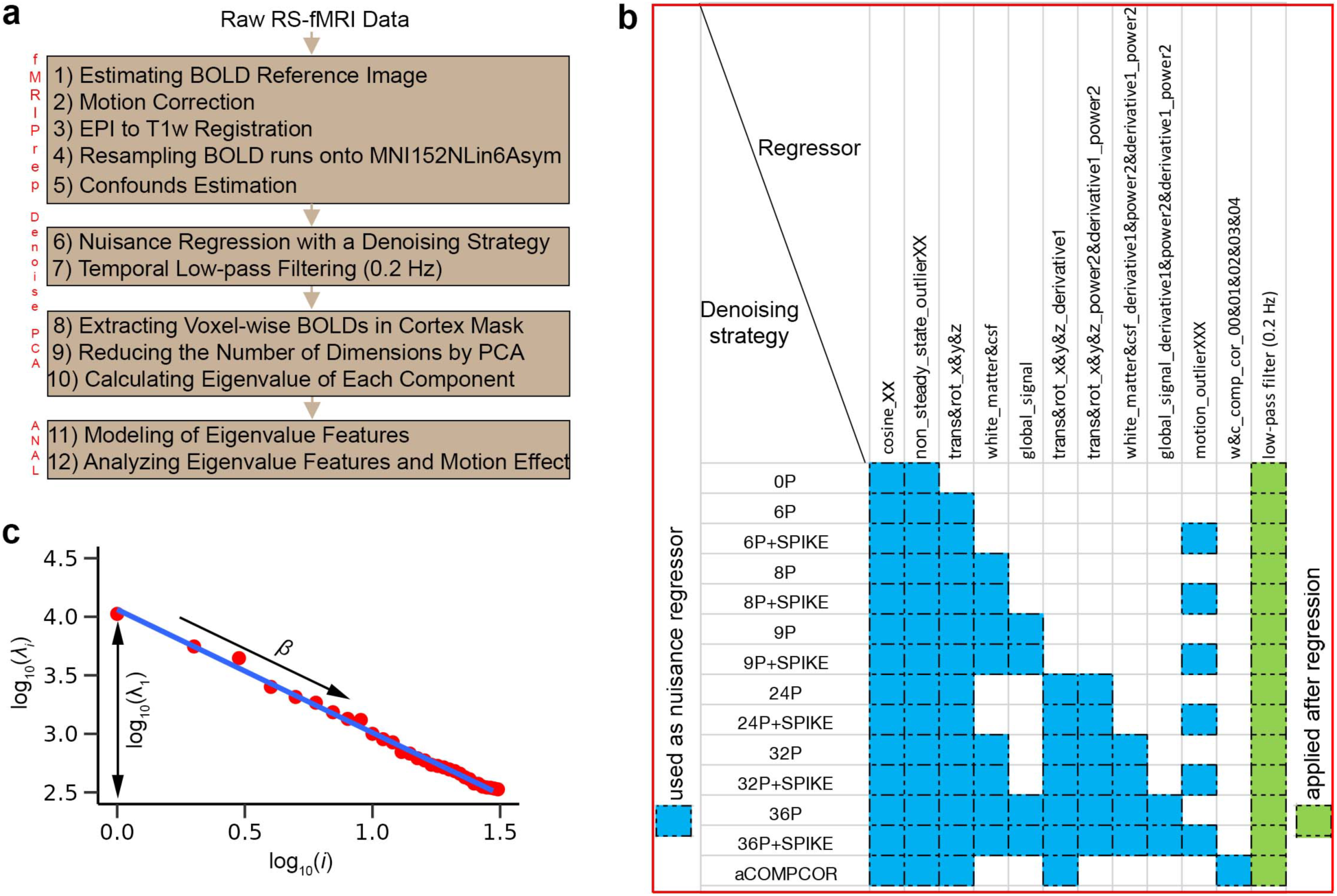
Flowchart of the study design, denoising strategies, and illustration of eigenvalue features. **a)** Overview of the study workflow. The flowchart consists of four main steps: (1) preprocessing RS-fMRI data using the fMRIPrep software; (2) applying various denoising strategies followed by temporal low-pass filtering with a cut-off frequency of 0.2 Hz; (3) extracting eigenvalues from cortical BOLD signals using principal component analysis (PCA); and (4) modeling eigenvalue-derived features and evaluating their properties and their relationship with the strategies and residual motion. **b)** Summary of the denoising strategies evaluated in this study. All regressors were derived from confound estimates produced by fMRIPrep. c) illustration of the derivation of the first eigenvalue (*λ*1) and slope of the eigenvalue spectrum b. *λi*, *λ*_1_, and *i* denote eigenvalue associated with component *i*, first eigenvalue, and component index, respectively.

This study utilizes two datasets. The first dataset, from a previous study (Yang, Vigotsky et al. 2022), includes 167 healthy controls (China167) (83 males, 84 females; mean age ± SD: 41 ± 14 years old). The second dataset comes from the Human Connectome Project 1200 (HCP1200) (Van Essen, Smith et al. 2013), which includes 926 young healthy volunteers after quality control (425 males, 501 females; mean age ± SD: 29 ± 4 years old).

### 2.2. Cortical Mask Generation

The cortical mask generation consisted of three steps and was consistent with the preceding preprocessing pipeline. First, for each dataset used in the study, subject-specific cortical region masks obtained after preprocessing and spatial normalization to MNI152 space were concatenated across subjects to form a four-dimensional (4D) image. Second, the 4D image was averaged across subjects and thresholded at 100% to generate a common cortical mask, ensuring that only voxels present in the cortical region of every subject were retained. Third, this common cortical mask was applied to the Harvard–Oxford cortical atlas (*HarvardOxford-cort-maxprob-thr25-2mm.nii.gz*) (Makris, Goldstein et al. 2006) to ensure that all voxels included in the cortical mask were confined within the MNI152 template space.

### 2.3. Confound Regressors and Denoising Strategies

Regressors were derived from the confound estimates generated by fMRIPrep and are illustrated in **Fig. 1b**. Each regressor type is described below:

**consine_XX**: Discrete cosine-basis regressors used as a temporal high-pass filter to remove low-frequency drift arising from physiological and/or scanner noise sources.

**non_steady_state_outlier_XX:** Indicator regressors for non-steady-state volumes, with a value of 1 for affected volumes and 0 elsewhere.

**trans&rot_x&y&z:** Six estimated head-motion parameters. All RS-fMRI volumes were realigned to the middle of the time series, yielding three translational (trans_x: left-right, trans_y: anterior-posterior, trans_z: superior-inferior) and three rotational parameters (rot_x: nodding “yes”, sagittal plane; rot_y: tilting left-right, frontal plane; rot_z: shaking “no”, transverse plane).

**white_matter&csf:** Mean signals extracted from anatomically derived, eroded white-matter (WM) and cerebrospinal fluid (CSF) masks.

**global_signal:** Mean signal extracted from the anatomically derived whole-brain mask

**trans&rot_x&y&z_derivative1:** First temporal derivatives of the six head-motion parameters

**trans&rot_x&y&z_power2&derivative1_power2:** Quadratic terms of the six motion parameters and their temporal derivatives.

**white_matter&csf_derivative1&power2&derivative1_power2:** First temporal derivatives and quadratic terms of the WM and CSF signals, as well as the temporal derivatives of those quadratic terms.

**global_signal_derivative1&power2&derivative1_power2:** First temporal derivative and quadratic term of the global signal, along with the temporal derivative of the quadratic term.

**motion_outlier_XXX:** Spike regressors used to censor volumes with excessive motion, defined as framewise displacement (FD) > 0.5 mm or standardized derivative of RMS variance over voxels (DVARS) > 1.5.

**w&c_comp_cor_00&01&02&03&04:** The first five principal components computed from WM and CSF masks using CompCor (Behzadi, Restom et al. 2007).

Fourteen denoising strategies were evaluated in this study, following approaches similar to those described in (Ciric, Wolf et al. 2017). Each strategy employed a different combination of confound regressors. As illustrated in **Fig. 2b**, the strategies were defined as follows:

- **0P**: cosine_XX + non_steady_state_outlier_XXX
- **6P**: 0P + trans&rot_x&y&z
- **6P+SPIKE** (**6PS**): 6P + motion_outlier_XXX
- **8P**: 6P + white_matter&csf
- **8P+SPIKE** (**8PS**): 8P + motion_outlier_XXX
- **9P**: 8P + global_signal
- **9P+SPIKE** (**9PS**): 9P + motion_outlier_XXX
- **24P**: 6P + trans&rot_x&y&z_derivative1 + trans&rot_x&y&z_power2&derivative1_power2
- **24P + SPIKE** (**24PS**): 24P + motion_outlier_XXX
- **32P**: 24P + white_matter&csf + white_matter&csf_derivative1&power2&derivative1_power2
- **32P + SPIKE** (**32PS**): 32P + motion_outlier_XXX
- **36P**: 32P + global_signal + global_signal_derivative1&power2&derivative1_power2
- **36P + SPKIE** (**36PS**): 36P + motion_outlier_XXX
- **aCOMPCOR** (**aC**): 6P + trans&rot_x&y&z_derivative1 + w&c_comp_cor_00&01&02&03&04

**Figure 2.**
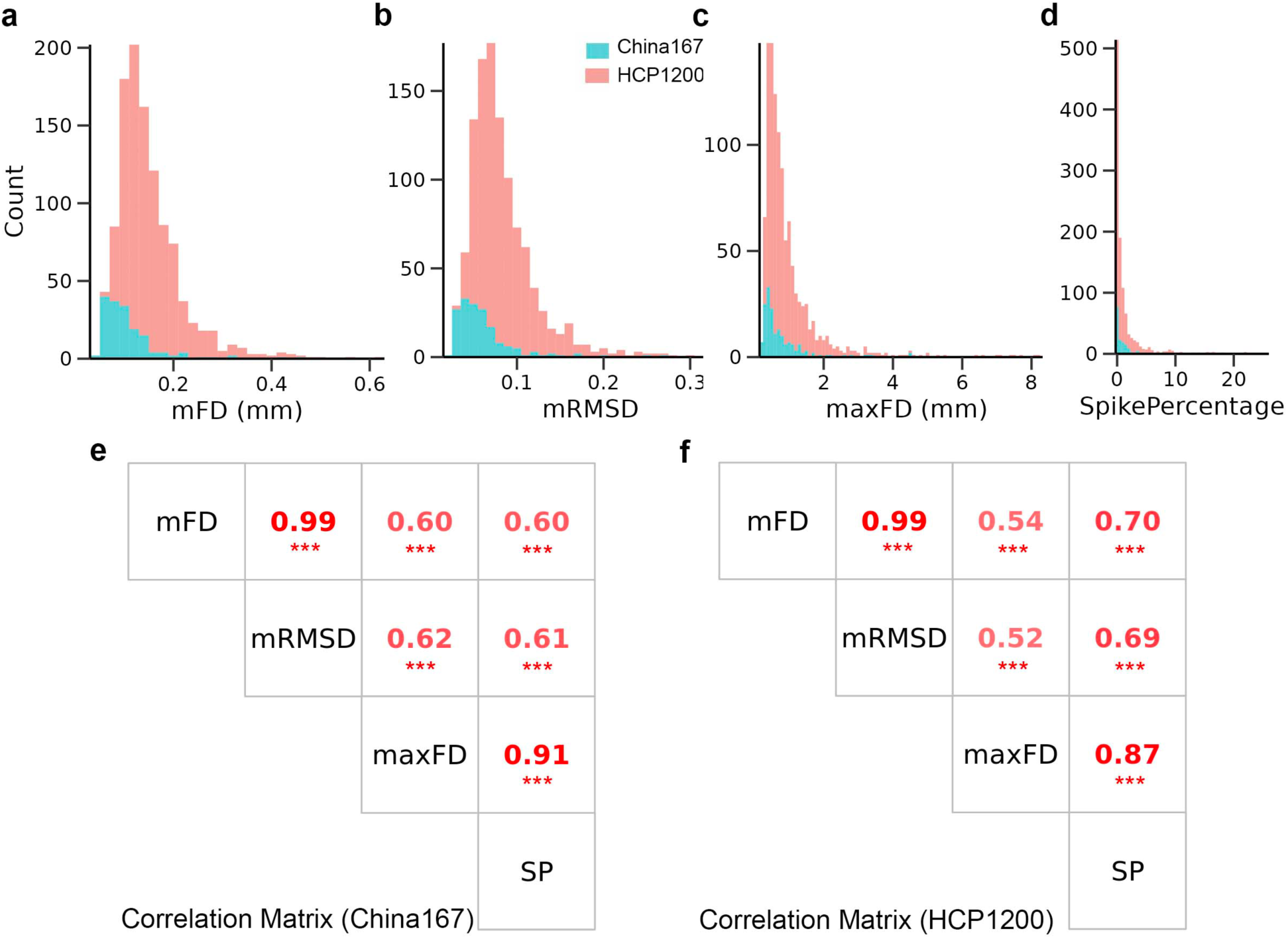
Histograms and correlation matrices of brain motion metrics for the China167 and HCP1200 datasets. **a)**, **b)**, **c)**, and **d)** display histograms of four motion metrics—mean framewise displacement (mFD), mean root mean square deviation (mRMSD), maximum framewise displacement (maxFD), and spike percentage (SP)—for the China167 (green) and HCP1200 (red) datasets. The HCP1200 dataset exhibits greater overall brain motion compared to the China167 dataset. **e)** and **f)** present the correlation matrices for the China167 and HCP1200 datasets, respectively. In both datasets, the four motion metrics are significantly correlated with one another. Notably, the correlation between mFD and mRMSD is nearly perfect (*r* = 0.99 in both datasets).

All 14 denoising strategies were followed by a fifth-order zero-phase Butterworth low-pass temporal filter with a cutoff frequency of 0.2 Hz. This filter yields a maximally flat amplitude response and a sharper transition between the passband and stopband in the frequency domain (Smith, Beckmann et al. 2013). The cutoff frequency was set to 0.2 Hz, rather than the more commonly used 0.08 or 0.1 Hz, to preserve higher-frequency components, as the denoised resting-state fMRI data were intended for subsequent calculation of the fractional amplitude of low-frequency fluctuations (fALFF) (Huang, Jabakhanji et al. 2026).

### 2.4. Linear Modeling of Eigenvalue Features to Capture Residual Motion Effects

As illustrated in **Fig. 1c**, the eigenspectrum of cortical BOLD signals exhibits an approximately linear relationship between log_10_-transformed eigenvalues and log_10_-transformed component indices. This relationship motivates the use of a two-parameter linear model to quantitatively characterize eigenvalue features that may reflect residual motion effects after denoising:

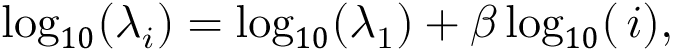

where *λ_i_* denotes the eigenvalue associated with component *i*, *λ*_1_ represents the first (largest) eigenvalue and serves as the intercept of the model. The parameter *β* captures the slope of the eigenspectrum and reflects the rate at which variance explained decreases across components. After model fitting, each subject under each denoising strategy is characterized by two eigenvalue feature parameters: log_10_(*λ*_1_) and *β*. These parameters provide a compact representation of the eigenspectrum and are used to assess residual motion in denoised RS-fMRI data.

To focus on the most informative components, only components satisfying log_10_(*i*) ≤ 1.5 (corresponding to approximately the first 40 components) were included in the analysis.

### 2.5. Relationship between two Eigenvalue Features

After extracting the two eigenvalue features, log_10_(*λ*_1_) and *β*, for each subject under 14 denoising strategies, the correlations between these features were computed across subjects for each denoising strategy within both the China167 and HCP1200 datasets. Results for four representative strategies (OP, 6P, 9P, and 36P) are reported because the other 10 strategies yielded highly similar results (log_10_( *λ*_1_) exhibited a strong and statistically significant association with *β*). Furthermore, for each dataset, the mean log_10_(*λ*_1_) and mean *β* (averaged across subjects for each denoising strategy) were first computed, and their correlation was then evaluated across the 14 denoising strategies. In addition, to assess the consistency of these two features, their correlations between the two datasets were also examined.

### 2.6. Relationship between Eigenvalue Features and Denoising Strategy

To illustrate how log_10_(*λ*_1_) (or *β*) is influenced by different denoising strategies, three subjects representing low, medium, and high motion levels—quantified by mFD (mean framewise displacement) and SP (spike percentage)—were selected from the China167 and HCP1200 datasets. For each subject, the relationship between log_10_(*λ_i_*) and log_10_(*i*) is depicted. Specifically, log_10_(*λ*_1_) is evaluated across four representative denoising strategies and their corresponding variants incorporating spike censoring. Additionally, group-level effects are examined by computing the mean log_10_(*λ*_1_) and mean *β*, enabling a comparison of how eigenvalue features vary across denoising approaches.

### 2.7. Relationship between log_10_(*λ*_1_) and Residual Motion

After establishing a strong association between log_10_(*λ*_1_) and *β* across subjects, denoising strategies, and datasets, only log_10_(*λ*_1_) is retained for subsequent analyses of its relationship with residual motion. For each strategy, we calculated the correlation between log_10_(*λ*_1_) and mFD across subjects to evaluate the relationship between log_10_(*λ*_1_) and residual motion within a strategy.

To further examine the relationship between residual motion and functional connectivity, we assessed the correlation between mean log_10_(*λ*_1_) and the percentage of edges “significantly” associated with motion (*p* < 0.05) across 14 denoising strategies in the China167 and HCP1200 datasets. The percentage of edges “significantly” associated with motion, also referred to as QC-

FC (quality control-functional connectivity) correlations, is widely used to evaluate the effectiveness of motion artifact removal in RS-fMRI studies (Power, Schlaggar et al. 2015, Ciric, Wolf et al. 2017), calculated as the proportion of connections (edges) between brain parcels (regions) whose Fisher’s *z* correlation coefficients are “statistically significantly” associated with mFD across subjects relative to the total number of possible connections in the network. In this study, both Power (Power, Cohen et al. 2011) and Schaefer (Schaefer, Kong et al. 2018) networks are used. Specifically, the percentage of motion-associated edges was defined as the proportion of functional connections (edges) between brain parcels (regions) whose Fisher’s *z* correlation coefficients were “significantly” associated with mFD across subjects, relative to the total number of possible connections in the network.

### 2.8. Relationship between Global Signal Regression or Spike Regression and Denoising Performance

After establishing that log_10_(*λ*_1_) reflects residual motion effects across RS-fMRI denoising strategies, we further examined how specific preprocessing choices influence this metric. Specifically, we constructed separate general linear models (GLMs) for the China167 and HCP1200 datasets to evaluate the effects of global signal regression (GSR) and spike regression on log_10_(*λ*_1_), while controlling for age and sex. For each dataset, variables were reshaped from a wide-format structure to a long-format structure across denoising strategies. In the resulting models, log_10_(*λ*_1_) was treated as the dependent variable, whereas the inclusion of GSR or spike regression was modeled as an explanatory variable using binary indicators denoting whether each procedure was applied within a given denoising strategy. Age and sex were included as covariates. The models were specified as:

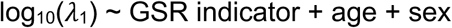

and

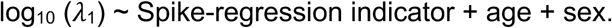

### 2.9. MRI scanning parameters

For the China167 dataset, subjects were scanned on a 3 Tesla GE-Discovery 750. T1 images were acquired with following parameters: voxel size = 1 × 1 × 1 mm^3^; TR/TE = 7.7/3.4 ms; flip angle = 12°; field of view = 256 × 256 mm^2^. RS-fMRI images were acquired with the following parameters: voxel size = 3.4375 × 3.4375 × 3.5 mm^3^; TR/TE = 2500/30 ms; flip angle = 90°; in-plane resolution = 64 × 64; field of view = 220 × 220 mm^2^; number of volumes = 230; slices per volume = 42, which covers the whole brain from the cerebellum to the vertex.

For the HCP1200 dataset, subjects were scanned on a 3 Tesla GE-Discovery 750. T1 images were acquired with following parameters: voxel size = 0.7 × 0.7 × 0.7 mm^3^; TR/TE = 2400/2.14 ms; flip angle = 8°; field of view = 224 × 224 mm^2^. RS-fMRI images were acquired with the following parameters: voxel size = 2 × 2 × 2 mm^3^; TR/TE = 720/33.1 ms; flip angle = 52°; field of view = 208 × 180 mm^2^; number of volumes = 1200; multiband accelerator = 8; slices per volume = 72, which covers the whole brain from the cerebellum to the vertex.

### 2.10. RS-fMRI data preprocessing using fMRIPrep software

The fMRI preprocessing is using fmriprep version of 23.1.4 (Esteban, Markiewicz et al. 2019), a Nipype-based tool (Gorgolewski, Burns et al. 2011, Gorgolewski KJ 2017). The details are described in (Huang, Jabakhanji et al. 2026).

## 3. Results

### 3.1. The four motion metrics are correlated with one another

As shown in **Fig. 2a-d**, histograms of the four motion metrics—mean framewise displacement (mFD), mean root mean square deviation (mRMSD), maximum framewise displacement (maxFD), and spike percentage (SP)—for the China167 (green) and HCP1200 (red) datasets indicate the HCP1200 dataset exhibits greater overall brain motion compared to the China167 dataset. The corresponding correlation matrices in **Fig. 2e-f** further demonstrate that the four motion metrics are strongly correlated with one another (all correlations *r* > 0.5 and *p* < 0.001). Notably, mFD and mRMSD show almost perfect association (*r* = 0.99, *p* < 0.001 in both datasets), while maxFD and SP are also highly correlated (*r* = 0.91 in China167 and *r* = 0.87 in HCP1200, *p* < 0.001 in both datasets). Based on these relationships, mFD and SP were selected as representative measures of motion in the present study.

### 3.2. **log_10_**( *λ*_1_) is strongly associated with *β*

As shown in **Fig.3**, for each of the four denoising strategies (0P, 6P, 9P, and 36P), log_10_(*λ*_1_) is negatively associated with *β* across subjects. The green open circles represent the mean values of log_10_(*λ*_1_) and *β*, which consistently shifted rightward and downward across the four denoising strategies in both datasets.

**Figure 3.**
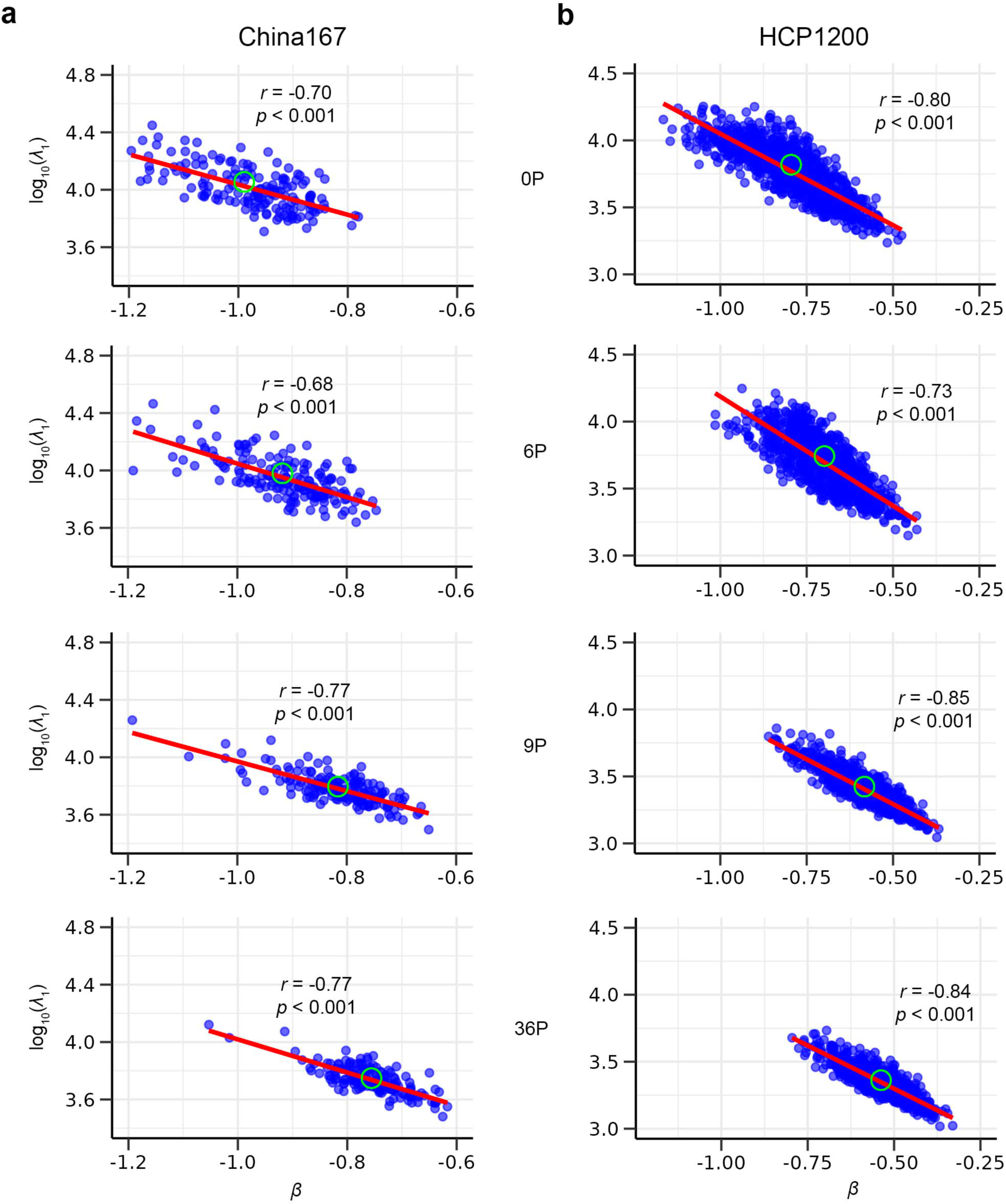
log_10_(*λ*_1_) is significantly associated with slope *β* in both the China167 and the HCP1200 datasets after applying four denoising strategies. a) Strong associations in the subjects from the China167 dataset, shown after regressing out 0P, 6P, 9P, and 36P confounds. **b)** Strong associations in the subjects from the HCP1200 dataset, shown after regressing out 0P, 6P, 9P, and 36P confounds. Note that the green empty circles indicate the mean position of log10(*λ*_1_) and *β*, which are moving right-down across the four denoising strategies for both the China167 and HCP1200 datasets.

Furthermore, as illustrated in **Fig. 4a** and **4b**, mean log_10_(*λ*_1_) remains strongly associated with mean *β* across 14 denoising strategies. This relationship is also highly consistent across datasets.

**Figure 4.**
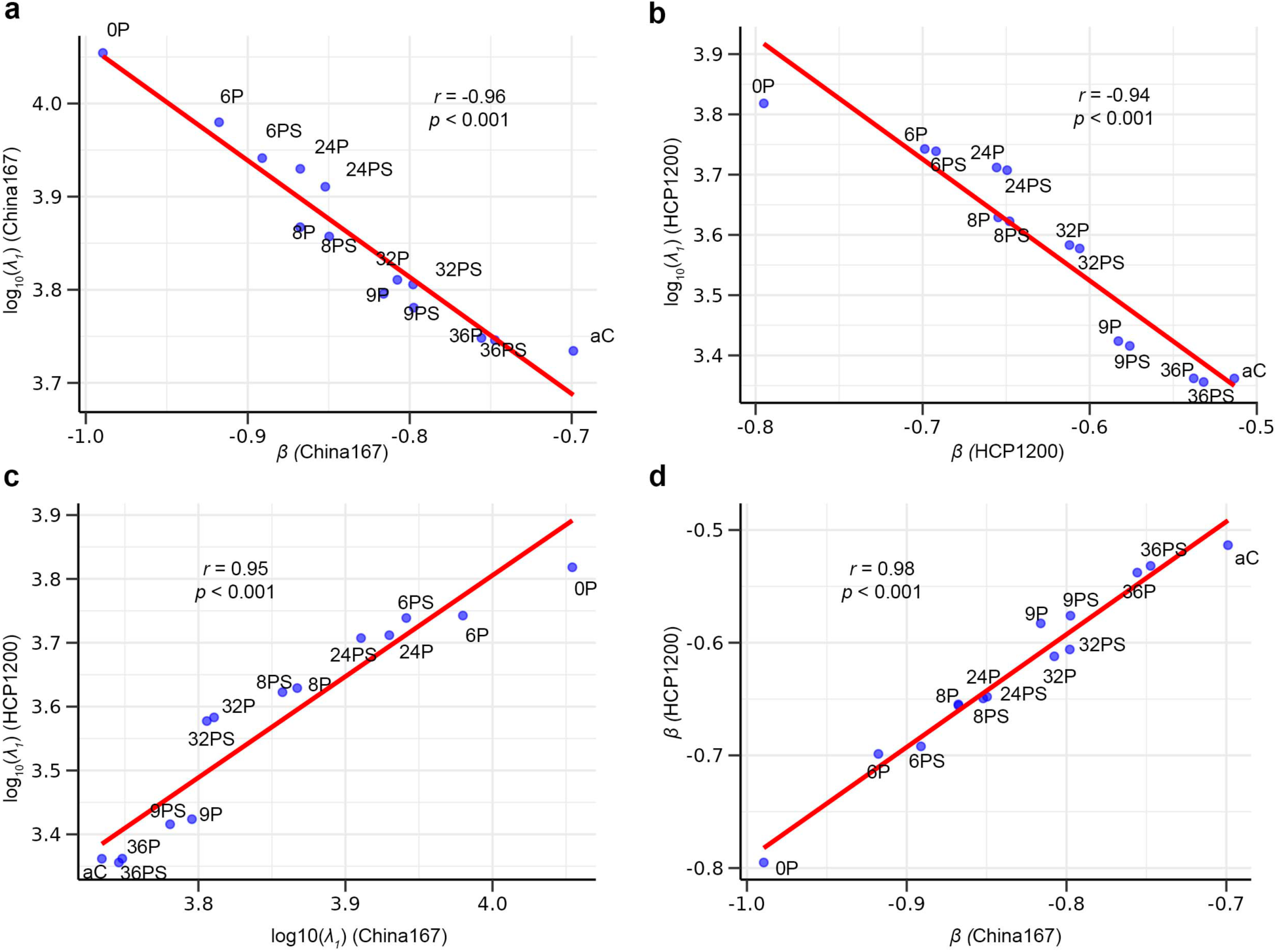
log_10_(*λ*_1_) is strongly associated with *β* within both the China167 and HCP1200 datasets across 14 denoising strategies. Moreover, both log_10_(*λ*_1_) and *β* show high correspondence between China167 and HCP1200 across the same 14 denoising strategies. a-b) Strong associations between log_10_(*λ*_1_) and *β* across 14 denoising strategies within the China167 and HCP1200 datasets. **c-d)** Strong associations of log_10_(*λ*_1_) and *β* across 14 denoising strategies between the China167 and HCP1200 datasets.

As shown in **Fig. 4c** and **4d**., there are strong cross-dataset associations for both mean log_10_(*λ*_1_) (*r* = 0.95; *p* < 0.001) and mean *β* (*r* = 0.98; *p* < 0.001) between the China167 and HCP1200 datasets.

Overall, these results demonstrate that log_10_(*λ*_1_) and *β* are robustly and consistently associated across subjects, denoising strategies, and datasets.

### 3.3. log_10_(*λ*_1_) and *β* are affected by denoising strategies

**Fig. 5** illustrates the effects of different denoising strategies on log_10_(*λ*_1_) and *β* across three motion levels in subjects from the China167 and HCP1200 datasets. At low motion (**Fig. 5a,b**), medium motion (**Fig. 5c,d**), and high motion (**Fig. 5e,f**), 0P, 6P, 9P, and 36P show a trend of decreasing log_10_(*λ*_1_) and increasing *β* in that order.

**Figure 5.**
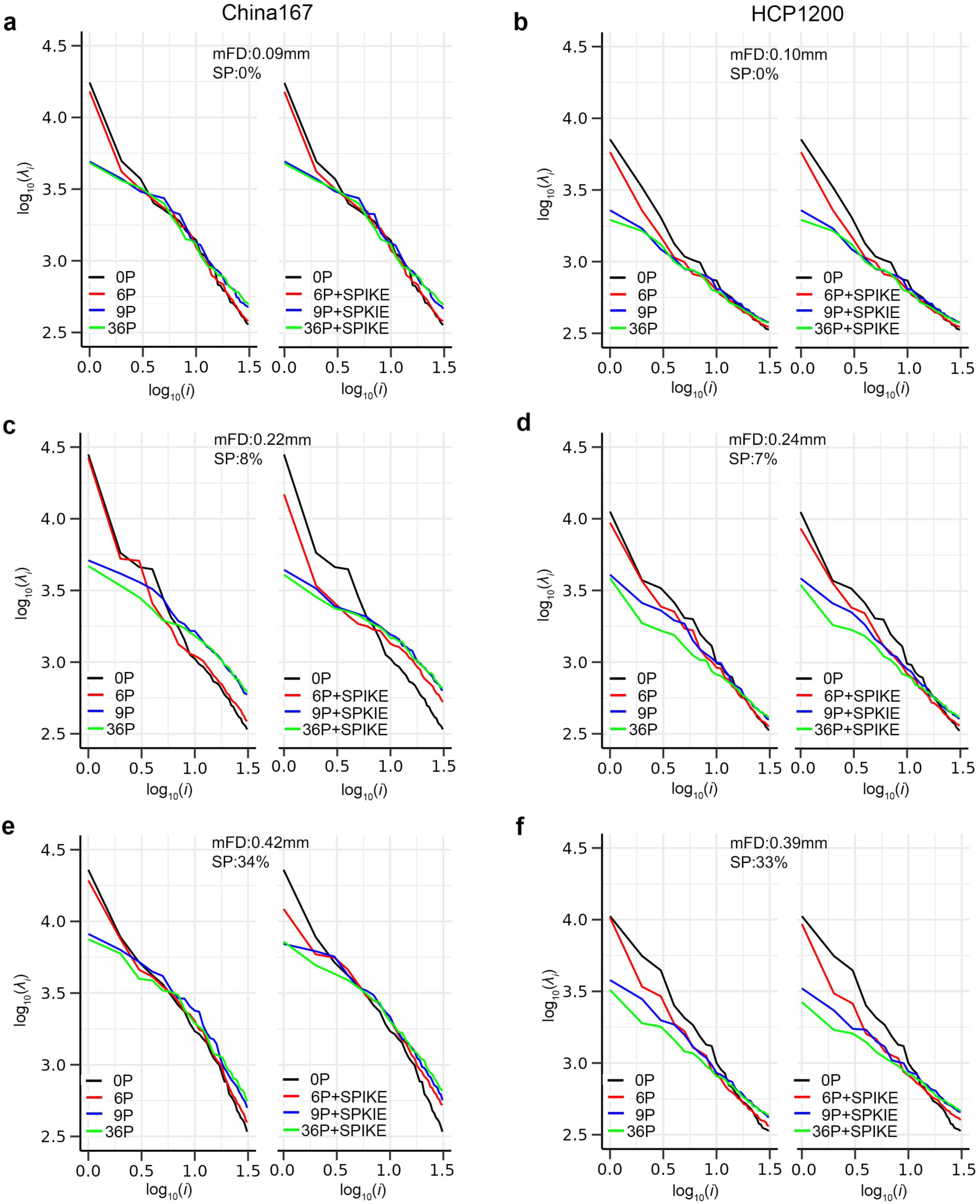
log_10_(*λ_i_*) plotted against log_10_(*i*) for three subjects exhibiting low, medium, and high motion from the China167 and HCP1200 datasets after applying four denoising strategies, each evaluated with and without spike regression. a) Low-motion subject from the China157 dataset, shown after regressing out 0P, 6P, 9P, and 36P confounds, displayed without spike regressors (left) and with spike regressors (right). **b)** Low-motion subject from the HCP1200 dataset. **c)** and **d)** Medium-motion subjects from the China167 and HCP1200 datasets. **e)** and **f)** High-motion subjects from the China157 and HCP1200 datasets. Note that 1) *λ_i_* and *i* denote eigenvalue associated with component *i* and component index, respectively; 2) mFD and SP refer to mean frame-wise displacement and spike percentage, respectively; 3) because SP = 0%, no spike regressors were included in **a**) and **b**).

Furthermore, except in the low-motion condition (SP = 0%), the spike-regression variants (6P+SPIKE, 9P+SPIKE, and 36P+SPIKE) exhibit slightly lower log_10_(*λ*_1_) and higher *β* compared with their corresponding base strategies (6P, 9P, and 36P), although these differences are minimal.

**Fig. 6** presents the mean log_10_(*λ*_1_) and *β* across subjects from the China167 and HCP1200 datasets after applying 14 different denoising strategies. Both subject-level and dataset-level results consistently indicate that the choice of denoising strategy influences log_10_(*λ*_1_) and *β*.

**Figure 6.**
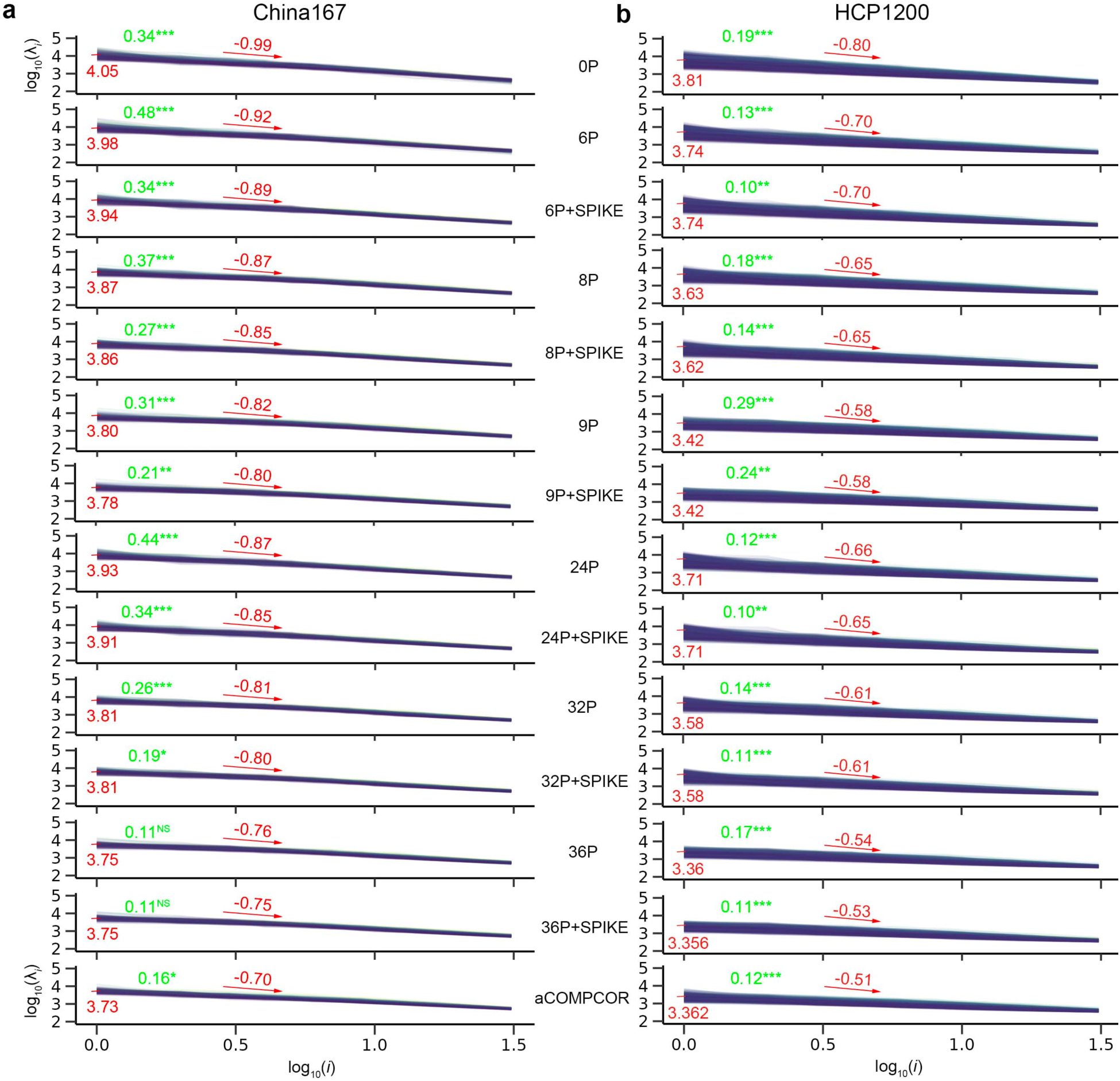
log_10_(*λ_i_*) plotted against log_10_(*i*) for subjects from the China167 and HCP1200 datasets after applying various denoising strategies with mean log_10_(*λ*_1_), mean *β*, and the correlation coefficient between log_10_(*λ*_1_) and mFD across subjects. a) China167 dataset. **b)** HCP1200 dataset. Red values denote the mean log_10_ *λ*_1_ and mean *β* for each denoising strategy. Green values denote the correlation coefficient between log_10_(*λ*_1_) and mFD across subjects for each strategy, where statistical significance is indicated as follows: *p* < 0.05 (*)*, p < 0.01* (**), *p* < 0.001 (***), and *p* > 0.05 (NS).

### 3.4. Association Between log10(*λ*_1_) and Residual Motion

Across subjects (within each denoising strategy), the correlation between log_10_(*λ*_1_) and mean framewise displacement (mFD) was computed (**Fig. 6**). Residual motion effects are reflected by positive correlations between log_10_(*λ*_1_) and mFD, indicating that a greater log_10_(*λ*_1_) reflects stronger residual motion effects. For the HCP1200 dataset, all denoising strategies produce log_10_(*λ*_1_) values positively associated with motion, indicating persistent motion-related effects after denoising. In contrast, for the China167 dataset, 12 out of the 14 strategies exhibit positive correlations (*p* < 0.05), except for the P36 and P36 + spike regression strategies.

Across denoising strategies, mean log_10_(*λ*_1_) strongly correlates with the QC-FC metric (**Fig. 7**). These edges were based on predefined functional network parcellations (Power and Schaefer); each edge represents connectivity between nodes within or across large-scale brain networks. Therefore, this proportion reflects how widely motion-related artifacts are distributed across functional networks. The observed relationship indicates that greater log_10_(*λ*_1_) values are associated with more widespread motion-related contamination affecting network-level connectivity patterns. This pattern is consistent across both the China167 (**Fig. 7a,c**) and HCP1200 (**Fig. 7b,d**) datasets and both network definitions when using both the Power and Schaefer networks.

**Figure 7.**
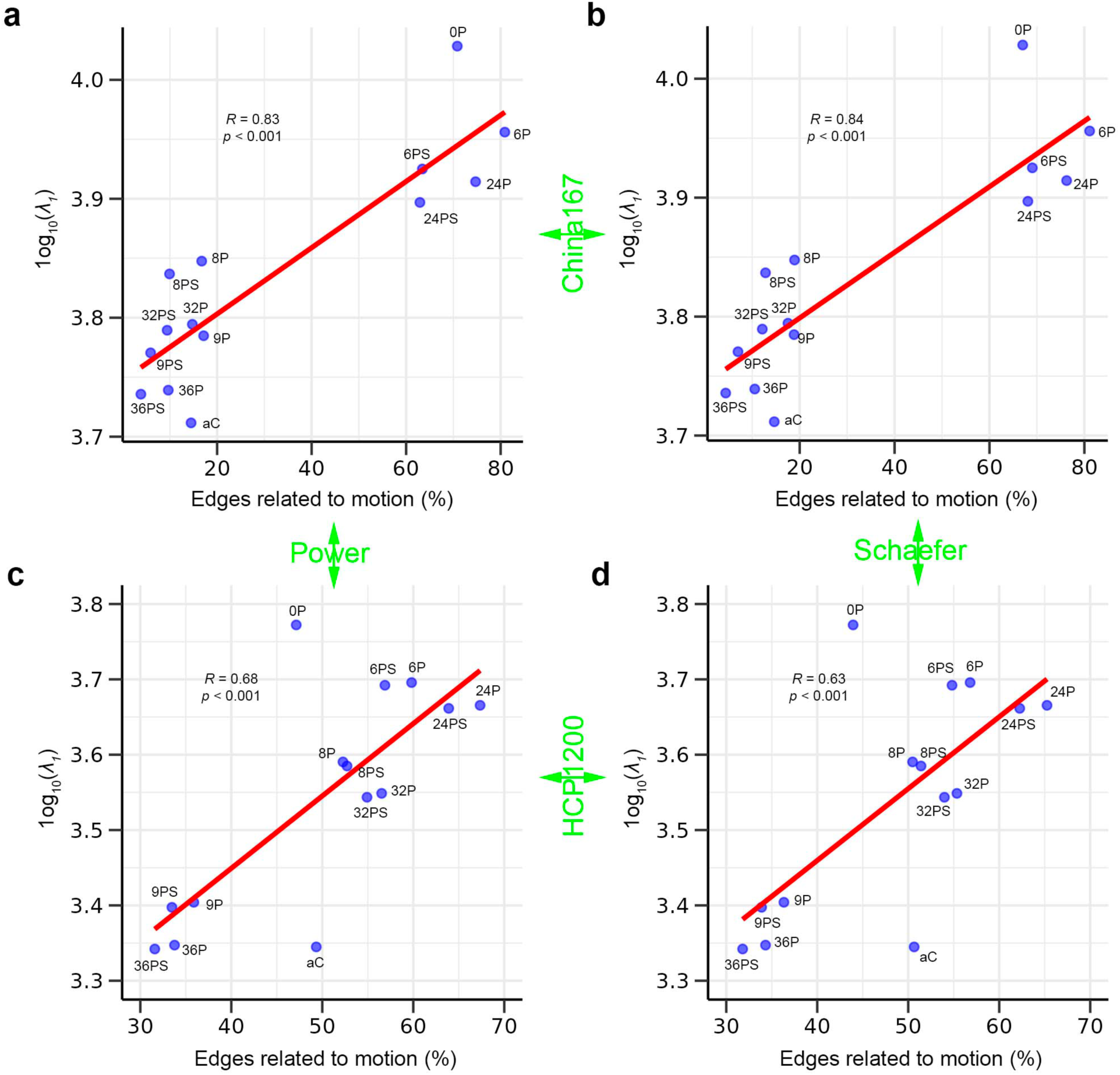
Mean log_10_(*λ*_1_) against the percentage of edges significantly associated with motion across 14 denoising strategies for subjects from the China167 and HCP1200 datasets on the Power and Schaefer networks. a) China167 dataset using the Power network. **b)** China167 dataset using the Schaefer network. **c)** HCP1200 dataset using the Power network. **d)** HCP1200 dataset using the Schaefer network. *λ*_1_ refers to the first eigenvalue; P, PS, and aC indicate physiological time series, physiological time series + spike regression and aCompCor, respectively.

Together, these results demonstrate a robust association between log_10_(*λ*_1_) and motion- related effects across both subject- and strategy-level analyses.

### 3.5. Global signal regression improves denoising performance, whereas spike regression shows dataset-dependent effects

As shown in **Fig. 8**, denoising strategies that include global signal regression yield consistently, though modestly, lower log_10_(*λ*_1_) than those without global signal regression in both the China167 and HCP1200 datasets. In contrast, spike regression has much more modest effects, which may be dataset-dependent. Specifically, strategies with spike regression yielded “significantly” lower first eigenvalues in the China167 dataset (*p* = 0.023) but not in the HCP1200 dataset (*p* = 0.212).

**Figure 8.**
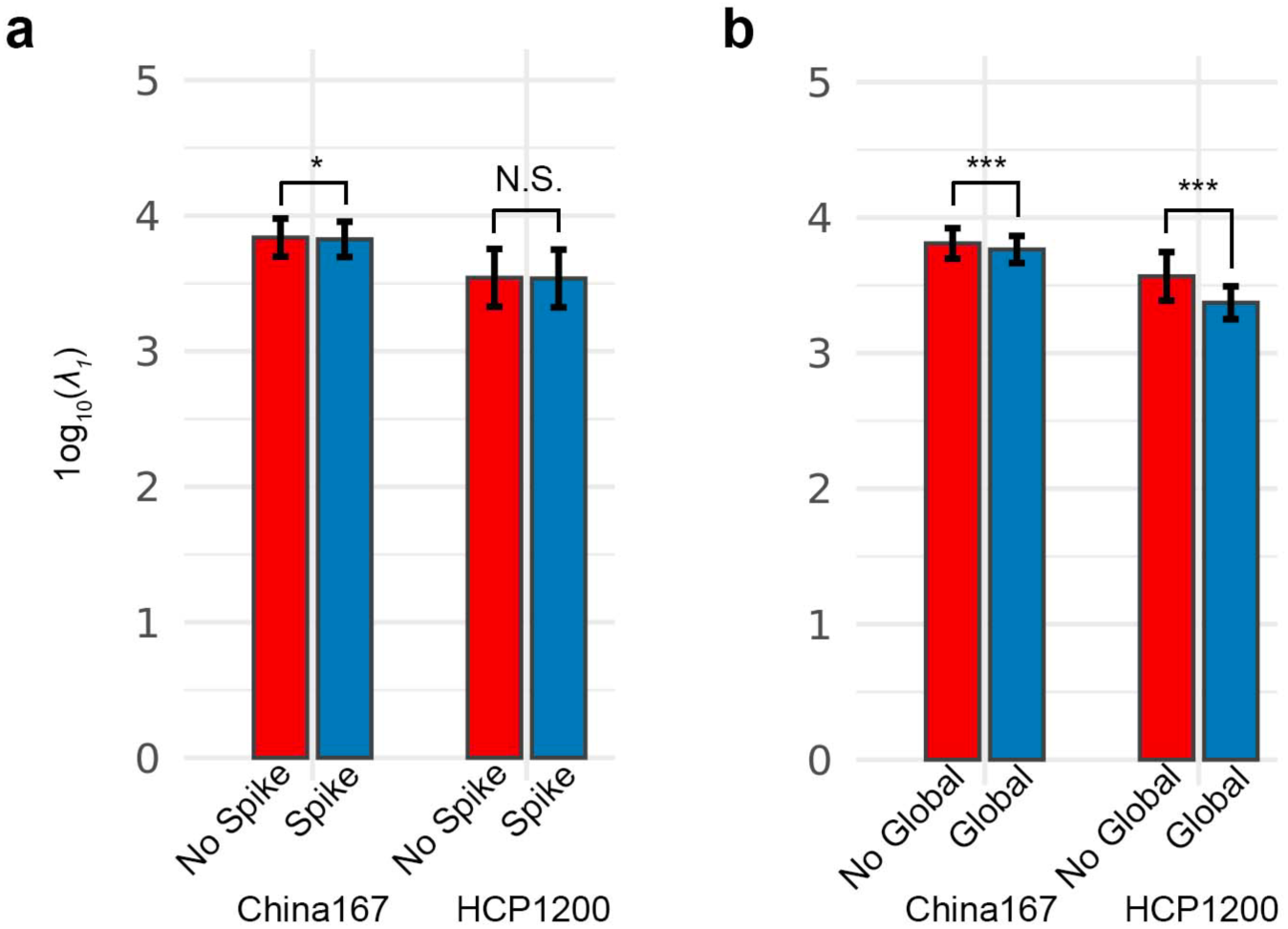
Effects of spike and global-signal regression on log_10_(*λ*_1_) in the China167 and HCP1200 datasets. **a)** Denoising strategies that include spike regression yield lower log_10_(*λ*_1_) compared with strategies without spike regression in the China167 dataset (left; *p* = 0.023), whereas the difference is “not significant” in the HCP1200 dataset (right; *p* = 0.212). **b)** Denoising strategies that include global-signal regression result in lower log_10_(*λ*_1_) than those without global-signal regression in both the China167 dataset (left; *p* < 0.001) and the HCP1200 dataset (right; *p* < 0.001). *λ*_1_ and N.S. refer to the first eigenvalue and non-significant (*p* > 0.05), respectively. Asterisks indicate significant thresholds: *p* < 0.05 (*) and *p* < 0.001 (***).

## Discussion

Eigenvalue analysis provides a compact representation of how variance is distributed across principal components. It has been widely applied in RS-fMRI analysis, including denoising, the characterization of functional connectivity, and structural mapping. In this study, we extend eigenvalue analysis to RS-fMRI quality control by introducing two features derived from the cortical eigenspectrum: log_10_(*λ*_1_), which reflects the relative dominance of the leading eigenvalue, and *β*, which characterizes the decay profile of the eigenspectrum. Notably, these two features are strongly correlated. This arises partly because eigenvalues are a zero-sum game: the original (*z*-scored) data has a fixed variance, so a greater first eigenvalue will be accompanied by lesser higher eigenvalues. (Notably, this is not mathematically guaranteed here since we do not use *all* components.) This strong association indicates that either feature can serve as a reliable and computationally efficient metric for group-level quality control after denoising.

Across denoising strategies, mean log_10_(*λ*_1_) is strongly positively correlated with the QC- FD correlations. This likely reflects spurious global or semi-global covariance patterns from motion effects. Further, this finding highlights the advantage of eigenspectrum analysis as a data-driven, model-free approach for detecting subtle artifacts that QC-FD may not adequately capture, partly because it is sensitive to how “significance” thresholds are implemented (Power, Barnes et al. 2012, Power, Mitra et al. 2014). In addition, because these features can be computed at the individual-subject level and averaged across subjects, they can be used for individual-subject denoising and facilitate straightforward comparisons across subjects and cohorts. These properties may induce a practical framework for subject-level quality control, in which inclusion or exclusion decisions for subsequent statistical analyses are guided by deviations in the mean log_10_(*λ*_1_) relative to a group reference distribution. This approach provides an objective and quantitative criterion, reduces reliance on arbitrary thresholds, and mitigates the influence of residual motion and noise-related artifacts.

In terms of denoising performance, the 36-parameter model with spike regression (36P + spike; 36PS) yields the lowest mean log₁₀(*λ*_1_), indicating the best performance for the HCP1200 dataset. This result is consistent with findings based on QC–FC correlations for the same dataset. However, for the China167 dataset, the aCompCor (aC) strategy achieves the lowest mean log₁₀(*λ*_1_), whereas 36PS still performs best in terms of QC–FC correlations. Notably, the difference in mean log₁₀(*λ*_1_) between 36PS and aC is small. Furthermore, the overall weak association between mean framewise displacement (mFD) and log₁₀(*λ*_1_) across subjects in both datasets suggests that 36PS is among the more robust denoising strategies evaluated. We also observed that the difference in mean log₁₀(*λ*_1_) between 9PS and 36PS is relatively small. Considering the preservation of degrees of freedom in downstream statistical analyses, 9PS may therefore represent a reasonable alternative. In contrast, although the 24PS model incorporates a larger set of motion-related regressors, it performs worse than 9PS on both log₁₀(*λ*_1_) and QC– FC correlation metrics. This observation is consistent with prior findings that increasing the number of motion regressors does not necessarily improve denoising and may instead remove variance that includes meaningful neural signal or structured fluctuations (Bright and Murphy 2015, Yang, Zhuang et al. 2019). By comparison, models such as 9PS that incorporate WM, CSF, and global signal regressors may capture variance beyond what motion regressors alone explain. Consistent with this interpretation, denoising strategies that include global signal regression show lower log₁₀(*λ*_1_) in both datasets (China167: *p* < 0.001; HCP1200: *p* < 0.001). In addition, the effect of spike regression appears to vary across datasets. Strategies including spike regression are associated with lower first eigenvalues in the China167 dataset (*p* = 0.023), but not in the HCP1200 dataset (*p* = 0.212). This pattern is broadly consistent with prior work showing that motion censoring (scrubbing) can improve data quality and reduce variance in some cases but may also have variable effects depending on dataset characteristics and the extent of motion contamination (Power, Barnes et al. 2012, Power, Mitra et al. 2014, Ciric, Wolf et al. 2017). Such variability may reflect differences in motion profiles, acquisition protocols, or preprocessing pipelines (Siegel, Power et al. 2014). Overall, these findings indicate that denoising performance depends not only on the number of regressors but also on their composition. Approaches that combine motion, physiological, and global components tend to perform relatively well in the present analyses, although the relative contribution of each component cannot be fully determined from these results alone.

## Limitations and Future Work

This study focuses on cortical BOLD signals and does not examine subcortical regions. However, subcortical structures are known to exhibit distinct BOLD signal characteristics compared to cortical areas (Drew 2019, Kim, Taylor et al. 2022), and incorporating these regions may provide additional insights. Second, as shown in **Fig. 7c–d**, the aCompCor denoising strategy demonstrates strong denoising performance in terms of log_10_(*λ*_1_) but relatively poor performance with respect to the QC–FD correlation metric. The underlying reasons for this discrepancy remain unclear and warrant further investigation.

## Conclusions

In this study, we introduce an eigenvalue-based framework that uses log_10_(*λ*_1_) and spectral slope *β* to compactly characterize RS-fMRI variance-covariance structure. These features are consistent across datasets and denoising strategies, systematically affected by denoising, and sensitive to residual motion. In particular, higher log_10_(*λ*_1_) reflects greater motion-related contamination and aligns with widespread motion–functional connectivity associations. Overall, eigenvalue signatures provide a simple, robust tool for evaluating denoising performance and assessing data quality in RS-fMRI. The corresponding script is publicly available at: https://github.com/lejianhuang/ALFF.

## Data available

The script is available in https://github.com/lejianhuang/EigenvalueSignature. The HCP1200 dataset is downloaded from the Human Connectome Project: https://www.humanconnectome.org/study/hcp-young-adult/. The China167 dataset will be available in https://openpain.org once upon the ongoing project is complete.

## Acknowledgement

We thank NIH (grant 1P50DA044121-01A1) for funding data analysis.

## Author Contributions

All authors contributed substantially to this work. A. V. A supervised the overall project. L. H. and A. D. V. conceived and designed the study and developed the methodology. L. H., and A. D. V. analyzed the data. L. H. drafted the initial manuscript. A. D. V. and A. V. A. revised the manuscript. All authors reviewed, revised, and approved the final version of the manuscript.

## Reference

Bansal, R. and B. S. Peterson (2021). “Use of random matrix theory in the discovery of resting state brain networks.” Magn Reson Imaging 77: 69–87.

Behzadi, Y., K. Restom, J. Liau and T. T. Liu (2007). “A component based noise correction method (CompCor) for BOLD and perfusion based fMRI.” Neuroimage 37(1): 90–101.

Bright, M. G. and K. Murphy (2015). “Is fMRI “noise” really noise? Resting state nuisance regressors remove variance with network structure.” Neuroimage 114: 158–169.

Ciric, R., D. H. Wolf, J. D. Power, D. R. Roalf, G. L. Baum, K. Ruparel, R. T. Shinohara, M. A. Elliott, S. B. Eickhoff, C. Davatzikos, R. C. Gur, R. E. Gur, D. S. Bassett and T. D. Satterthwaite (2017). “Benchmarking of participant-level confound regression strategies for the control of motion artifact in studies of functional connectivity.” Neuroimage 154: 174–187.

Drew, P. J. (2019). “Vascular and neural basis of the BOLD signal.” Curr Opin Neurobiol 58: 61–69.

Esteban, O., C. J. Markiewicz, R. W. Blair, C. A. Moodie, A. I. Isik, A. Erramuzpe, J. D. Kent, M. Goncalves, E. DuPre, M. Snyder, H. Oya, S. S. Ghosh, J. Wright, J. Durnez, R. A. Poldrack and K. J. Gorgolewski (2019). “fMRIPrep: a robust preprocessing pipeline for functional MRI.” Nat Methods 16(1): 111–116.

Friston, K. J., S. Williams, R. Howard, R. S. Frackowiak and R. Turner (1996). “Movement-related effects in fMRI time-series.” Magn Reson Med 35(3): 346–355.

Ghosh, S., A. Raj and S. Nagarajan (2024). “A Joint Subspace Mapping Between Structural and Functional Brain Connectomes.” Journal of Computational Neuroscience 52: S94–S94.

Golestani, A. M. and J. J. Chen (2022). “Performance of Temporal and Spatial Independent Component Analysis in Identifying and Removing Low-Frequency Physiological and Motion Effects in Resting-State fMRI.” Front Neurosci 16: 867243.

Gorgolewski, K., C. D. Burns, C. Madison, D. Clark, Y. O. Halchenko, M. L. Waskom and S. S. Ghosh (2011). “Nipype: a flexible, lightweight and extensible neuroimaging data processing framework in python.” Front Neuroinform 5: 13.

Gorgolewski KJ, E. O., Ellis DG, Notter MP, Ziegler E, Johnson H, Hamalainen C, Yvernault B, Burns C, Manhães-Savio A, Jarecka D, Markiewicz CJ, Salo T, Clark D, Waskom M, Wong J, Modat M, Dewey BE, Clark MG, Dayan M, Loney F, Madison C, Gramfort A, Keshavan A, Berleant S, Pinsard B, Goncalves M, Clark D, Cipollini B, Varoquaux G, Wassermann D, Rokem A, Halchenko YO, Forbes J, Moloney B, Malone IB, Hanke M, Mordom D, Buchanan C, Pauli WM, Huntenburg JM, Horea C, Schwartz Y, Tungaraza R, Iqbal S, Kleesiek J, Sikka S, Frohlich C, Kent J, Perez-Guevara M, Watanabe A, Welch D, Cumba C, Ginsburg D, Eshaghi A, Kastman E, Bougacha S, Blair R, Acland B, Gillman A, Schaefer A, Nichols BN, Giavasis S, Erickson D, Correa C, Ghayoor A, Küttner R, Haselgrove C, Zhou D, Craddock RC, Haehn D, Lampe L, Millman J, Lai J, Renfro M, Liu S, Stadler J, Glatard T, Kahn AE, Kong X-Z, Triplett W, Park A, McDermottroe C, Hallquist M, Poldrack R, Perkins LN, Noel M, Gerhard S, Salvatore J, Mertz F, Broderick W, Inati S, Hinds O, Brett M, Durnez J, Tambini A, Rothmei S, Andberg SK, Cooper G, Marina A, Mattfeld A, Urchs S, Sharp P, Matsubara K, Geisler D, Cheung B, Floren A, Nickson T, Pannetier N, Weinstein A, Dubois M, Arias J, Tarbert C, Schlamp K, Jordan K, Liem F, Saase V, Harms R, Khanuja R, Podranski K, Flandin G, Papadopoulos Orfanos D, Schwabacher I, McNamee D, Falkiewicz M, Pellman J, Linkersdörfer J, Varada J, Pérez-García F, Davison A, Shachnev D, Ghosh S. (2017). Nipype: a flexible, lightweight and extensible neuroimaging data processing framework in Python.

Huang, L., R. Jabakhanji, A. D. Vigotsky, P. Branco, M. N. Baliki and A. V. Apkarian (2026). “Revisiting Amplitude of Low-Frequency Fluctuations (ALFF) in Resting-State fMRI: Clarifications and Improvements.” Hum Brain Mapp 47(5): e70506.

Huang, S., A. D. Vigotsky, A. V. Apkarian and L. Huang (2024). “Body mass index associated with respiration predicts motion in resting-state functional magnetic resonance imaging scans.” Hum Brain Mapp 45(13): e70015.

Jolliffe, I. T. (2002). Principal component analysis. New York, Springer.

Kim, J. H., A. J. Taylor, M. Himmelbach, G. E. Hagberg, K. Scheffler and D. Ress (2022). “Characterization of the blood oxygen level dependent hemodynamic response function in human subcortical regions with high spatiotemporal resolution.” Front Neurosci 16: 1009295.

Leonardi, N., J. Richiardi, M. Gschwind, S. Simioni, J. M. Annoni, M. Schluep, P. Vuilleumier and D. Van De Ville (2013). “Principal components of functional connectivity: a new approach to study dynamic brain connectivity during rest.” Neuroimage 83: 937–950.

Makris, N., J. M. Goldstein, D. Kennedy, S. M. Hodge, V. S. Caviness, S. V. Faraone, M. T. Tsuang and L. J. Seidman (2006). “Decreased volume of left and total anterior insular lobule in schizophrenia.” Schizophr Res 83(2-3): 155–171.

Pavlovich, K., J. Pang, A. Holmes, T. Constable and A. Fornito (2025). “The efficacy of resting-state fMRI denoising pipelines for motion correction and behavioural prediction.” Imaging Neurosci (Camb) 3.

Power, J. D., K. A. Barnes, A. Z. Snyder, B. L. Schlaggar and S. E. Petersen (2012). “Spurious but systematic correlations in functional connectivity MRI networks arise from subject motion.” Neuroimage 59(3): 2142–2154.

Power, J. D., A. L. Cohen, S. M. Nelson, G. S. Wig, K. A. Barnes, J. A. Church, A. C. Vogel, T. O. Laumann, F. M. Miezin, B. L. Schlaggar and S. E. Petersen (2011). “Functional network organization of the human brain.” Neuron 72(4): 665–678.

Power, J. D., A. Mitra, T. O. Laumann, A. Z. Snyder, B. L. Schlaggar and S. E. Petersen (2014). “Methods to detect, characterize, and remove motion artifact in resting state fMRI.” Neuroimage 84: 320–341.

Power, J. D., B. L. Schlaggar and S. E. Petersen (2015). “Recent progress and outstanding issues in motion correction in resting state fMRI.” Neuroimage 105: 536–551.

Pruim, R. H. R., M. Mennes, D. van Rooij, A. Llera, J. K. Buitelaar and C. F. Beckmann (2015). “ICA-AROMA: A robust ICA-based strategy for removing motion artifacts from fMRI data.” Neuroimage 112: 267–277.

Reddy, N. A., K. M. Zvolanek, S. Moia, C. Caballero-Gaudes and M. G. Bright (2024). “Denoising task-correlated head motion from motor-task fMRI data with multi-echo ICA.” Imaging Neurosci (Camb) 2.

Satterthwaite, T. D., D. H. Wolf, J. Loughead, K. Ruparel, M. A. Elliott, H. Hakonarson, R. C. Gur and R. E. Gur (2012). “Impact of in-scanner head motion on multiple measures of functional connectivity: relevance for studies of neurodevelopment in youth.” Neuroimage 60(1): 623–632.

Schaefer, A., R. Kong, E. M. Gordon, T. O. Laumann, X. N. Zuo, A. J. Holmes, S. B. Eickhoff and B. T. T. Yeo (2018). “Local-Global Parcellation of the Human Cerebral Cortex from Intrinsic Functional Connectivity MRI.” Cereb Cortex 28(9): 3095–3114.

Siegel, J. S., J. D. Power, J. W. Dubis, A. C. Vogel, J. A. Church, B. L. Schlaggar and S. E. Petersen (2014). “Statistical improvements in functional magnetic resonance imaging analyses produced by censoring high-motion data points.” Hum Brain Mapp 35(5): 1981–1996.

Smith, S. M., C. F. Beckmann, J. Andersson, E. J. Auerbach, J. Bijsterbosch, G. Douaud, E. Duff, D. A. Feinberg, L. Griffanti, M. P. Harms, M. Kelly, T. Laumann, K. L. Miller, S. Moeller, S. Petersen, J. Power, G. Salimi-Khorshidi, A. Z. Snyder, A. T. Vu, M. W. Woolrich, J. Xu, E. Yacoub, K. Ugurbil, D. C. Van Essen, M. F. Glasser and W. U.-M. H. Consortium (2013). “Resting-state fMRI in the Human Connectome Project.” Neuroimage 80: 144–168.

Soltysik, D. A., D. Thomasson, S. Rajan and N. Biassou (2015). “Improving the use of principal component analysis to reduce physiological noise and motion artifacts to increase the sensitivity of task-based fMRI.” J Neurosci Methods 241: 18–29.

Van Dijk, K. R., M. R. Sabuncu and R. L. Buckner (2012). “The influence of head motion on intrinsic functional connectivity MRI.” Neuroimage 59(1): 431–438.

Van Essen, D. C., S. M. Smith, D. M. Barch, T. E. Behrens, E. Yacoub, K. Ugurbil and W. U.-M. H. Consortium (2013). “The WU-Minn Human Connectome Project: an overview.” Neuroimage 80: 62–79.

Wang, M. B., J. P. Owen, P. Mukherjee and A. Raj (2017). “Brain network eigenmodes provide a robust and compact representation of the structural connectome in health and disease.” PLoS Comput Biol 13(6): e1005550.

Yang, L., A. D. Vigotsky, B. Wu, B. Shen, Z. Yan, A. V. Apkarian and L. Huang (2022). “Morphometric similarity networks discriminate patients with lumbar disc herniation from healthy controls and predict pain intensity.” Front Netw Physiol 2: 992662.

Yang, L., B. Wu, L. Fan, S. Huang, A. D. Vigotsky, M. N. Baliki, Z. Yan, A. V. Apkarian and L. Huang (2021). “Dissimilarity of functional connectivity uncovers the influence of participant’s motion in functional magnetic resonance imaging studies.” Hum Brain Mapp 42(3): 713–723.

Yang, Z., X. Zhuang, K. Sreenivasan, V. Mishra, D. Cordes and I. Alzheimer’s Disease Neuroimaging (2019). “Robust Motion Regression of Resting-State Data Using a Convolutional Neural Network Model.” Front Neurosci 13: 169.

Zajac, K. and J. Piersa (2013). “Eigenvalue Spectra of Functional Networks in fMRI Data and Artificial Models.” Artificial Intelligence and Soft Computing, Pt I 7894: 205–214.

Zhu, W., X. Ma, X. H. Zhu, K. Ugurbil, W. Chen and X. Wu (2022). “Denoise Functional Magnetic Resonance Imaging With Random Matrix Theory Based Principal Component Analysis.” IEEE Trans Biomed Eng 69(11): 3377–3388.

